# DomainFit: Identification of Protein Domains in cryo-EM maps at Intermediate Resolution using AlphaFold2-predicted Models

**DOI:** 10.1101/2023.11.28.569001

**Authors:** Jerry Gao, Max Tong, Chinkyu Lee, Jacek Gaertig, Thibault Legal, Khanh Huy Bui

## Abstract

Cryo-electron microscopy (cryo-EM) has revolutionized our understanding of macromolecular complexes, enabling high-resolution structure determination. With the paradigm shift to *in situ* structural biology recently driven by the ground-breaking development of cryo-focused ion beam milling and cryo-electron tomography, there are an increasing number of structures at sub-nanometer resolution of complexes solved directly within their cellular environment. These cellular complexes often contain unidentified proteins, related to different cellular states or processes. Identifying proteins at resolutions lower than 4 Å remains challenging because the side chains cannot be visualized reliably. Here, we present DomainFit, a program for automated domain-level protein identification from cryo-EM maps at resolutions lower than 4 Å. By fitting domains from artificial intelligence-predicted models such as AlphaFold2-predicted models into cryo-EM maps, the program performs statistical analyses and attempts to identify the proteins forming the density. Using DomainFit, we identified two microtubule inner proteins, one of them, a CCDC81 domain-containing protein, is exclusively localized in the proximal region of the doublet microtubule from the ciliate *Tetrahymena thermophila*. The flexibility and capability of DomainFit makes it a valuable tool for analyzing *in situ* structures.

## Introduction

In the last decade, cryo-electron microscopy (cryo-EM) has become a powerful technique to determine the structures of macromolecular complexes. With advances in cryo-EM image processing algorithms, large endogenous complexes have been solved at high resolutions such as the phycobilisomes in red algae 1, the mitochondrial membrane bending supercomplex 2 and the doublet microtubules of the cilia 3,4. Recently, *in situ* cryo-electron tomography (cryo-ET) from thin lamellae of cells prepared by a cryo-focused ion beam (cryo-FIB) instrument has emerged as a transformative technique, revolutionizing our understanding of cellular structures and molecular processes 5. With high-throughput tilt series acquisition and improvement of subtomogram averaging software 6–13, subtomogram averaging of complexes can reach sub-nanometer resolution and in some cases better than 4 Å resolution 11,14–16. Structures of protein complexes obtained *in situ* often contain unknown components, unlike *in vitro* reconstituted protein complexes. In addition, depending on the cellular context, space-and/or time-specific interactions with unknown proteins might be revealed by subtomogram averaging. Due to the lower resolution of these *in situ* structures, identification of proteins requires a different approach than traditional molecular modelling programs.

For high-resolution structures, there are many methods that can be used for the modelling and identification of unknown proteins including CryoID, DeeptracerID, FindMySequence, and modelAngelo 17–20. These programs trace and model the backbone of the protein density. Then, the identity of the protein is predicted by comparing the side-chain densities against a database of protein sequences. This approach was successful in determining many proteins in the doublet microtubule 21,22, radial spokes and central apparatus and the phycobilisomes 17. However, all these methods only work reliably at a resolution better than 4 Å.

In most cases, the attainable resolution by subtomogram averaging is lower than 4 Å. This is due to the low throughput of cryo-FIB milling and low abundance of the target protein complex. Furthermore, *in situ* structures can have problems related to orientation, sample thickness and signal-to-noise ratio. Therefore, it is a challenge to identify proteins from *in situ* subtomogram averages.

In the last few years, there has been a breakthrough in artificial intelligence (AI)-assisted protein structure prediction. Programs like AlphaFold 23, ColabFold 24 and RoseTTAfold 25 can produce accurate structure predictions. With the establishment of the AlphaFold Protein Structure Database (AlphaFold DB), everyone now has access to over 200 million predicted models. For certain organisms such as humans, mice, zebrafish and nematodes, AlphaFold2-predicted models of the entire proteome are available.

With the good accuracy of the AI-predicted models, it is possible to identify the fold signature of a protein at a resolution of less than 4 Å by fitting the AI-predicted models into the density. Using complementary data such as *in situ* chemical cross-linking mass spectrometry, quantitative mass spectrometry and/or BioID, it is possible to identify proteins from the fit list. For example, microtubule inner proteins (MIPs) were identified from the *in situ* subtomogram average of human sperm doublet microtubules at ∼6-7 Å using Colores Situs program by fitting over 21,000 AlphaFold2-predicted protein models of the mouse proteome 26. In another study, 38 proteins were identified from the mouse sperm by manually fitting AlphaFold2-predicted models of proteins found in the mass spectrometry analysis of the same sample 16.

While those studies illustrate that it is possible to identify well-structured proteins from a map lower than 4 Å resolution, the methods are not automated or are difficult to use. In addition, the criteria to evaluate and identify proteins from the fitting solutions are not clear from these studies. While the accuracy of the AI-predicted model is high for compact domains, the tertiary structure of the predicted model might not be correct, due in part to poor prediction of flexible regions. Therefore, protein identification is easier when domains are extracted from each AI-predicted model and used instead of the entire model.

In this work, we developed a program called DomainFit to identify domains of proteins from cryo-EM maps of lower resolution than 4 Å. DomainFit uses popular programs such as Phenix 27, R 28 and UCSF ChimeraX (ChimeraX) 29, making it easy to install and accessible to people. In addition, DomainFit is flexible and has a clear statistical approach to evaluate the identity of proteins. Lastly, DomainFit uses ChimeraX, which allows easy visualization to aid in evaluating the fit. Our workflow will be useful for the upcoming wave of *in situ* structures as cryo-FIB and cryo-ET become more popular.

## Results

### An automated pipeline for domain parsing and fitting into cryo-EM map

Overall, the program is designed to first download a database of AI-predicted models of candidate proteins (i.e. full species proteome or a limited list based on mass spectrometry studies), then divide each model into different domains and fit each individual domain into a segmented cryo-EM density. Finally, the program generates a statistical evaluation of all the fitted domains (Fig. 1A). The identification of the correct domain can be based purely on statistical analysis but also on other complementary data such as surrounding densities, protein size, chemical cross-linking and quantitative mass spectrometry.

**Figure 1:**
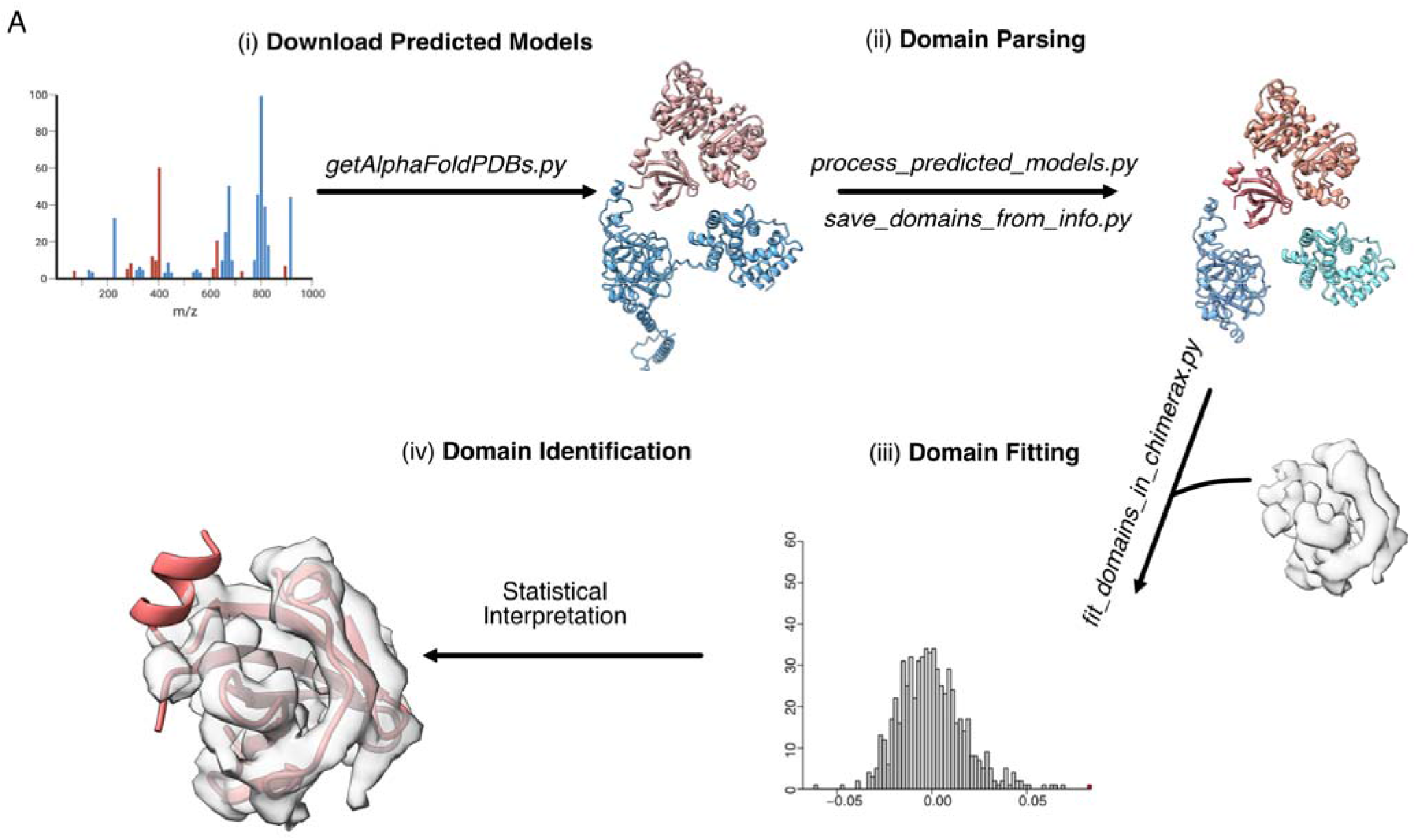
Workflow of DomainFit. (A) Overview of the program. (i) Download the predicted PDBs for all candidate proteins. (ii) Predict domains and split them into single PDB files. This step can be done through the script *process_predicted_models.py* or results from other programs such as DPAM. (iii) Fit each domain into the unknown density using ChimeraX nd generate the best fitting position of each domain and fitting scores. (iv) Interpret the fitting results based on p-value, correlation, and size of proteins.

The list of candidate proteins can be obtained from the mass spectrometry results of the target sample such as the proteome of isolated organelles or pull-down analysis. In case a targeted list is not available, it is possible to use AI-predicted models of the entire proteome of the organism, even though it will increase computational time significantly. In DomainFit, we provide a script *getAlphaFoldPDBs.py* to automatically download the AlphaFold2-predicted models from the AlphaFold DB using a list of protein UniProtIDs (Fig. 1A). The downside of downloading automatically from the AlphaFold DB is that for many organisms, high molecular weight proteins (> 100kDa) are not yet available in the database. These missing proteins can be predicted using ColabFold, RoseTTAFold or a local instance of AlphaFold.

Many ideas have been proposed to break PDB models into compact domains automatically 27,30,31. With AI-predicted models, it is possible to partition into domains based on the predicted alignment errors (PAE) of each model (Fig. 2A-D). PAE is the expected positional error for residue x if the predicted and actual models are aligned at residue y 23. Using an image-based approach, it is possible to partition predicted models into domains based on their respective PAE files (Fig. 2B-D) 30. We used *phenix.process_predicted_model* program from the Phenix package 27 for the partitioning of the PDB into domains because of its flexibility. *phenix.process_predicted_model* can function based on PAE but also based on the 3D arrangement of the proteins without PAE information. Therefore, we implemented a Python wrapper *process_predicted_models.py* to partition PDBs into domains in batch (Fig. 1A). Users can customize the options for parsing based on their needs.

**Figure 2.**
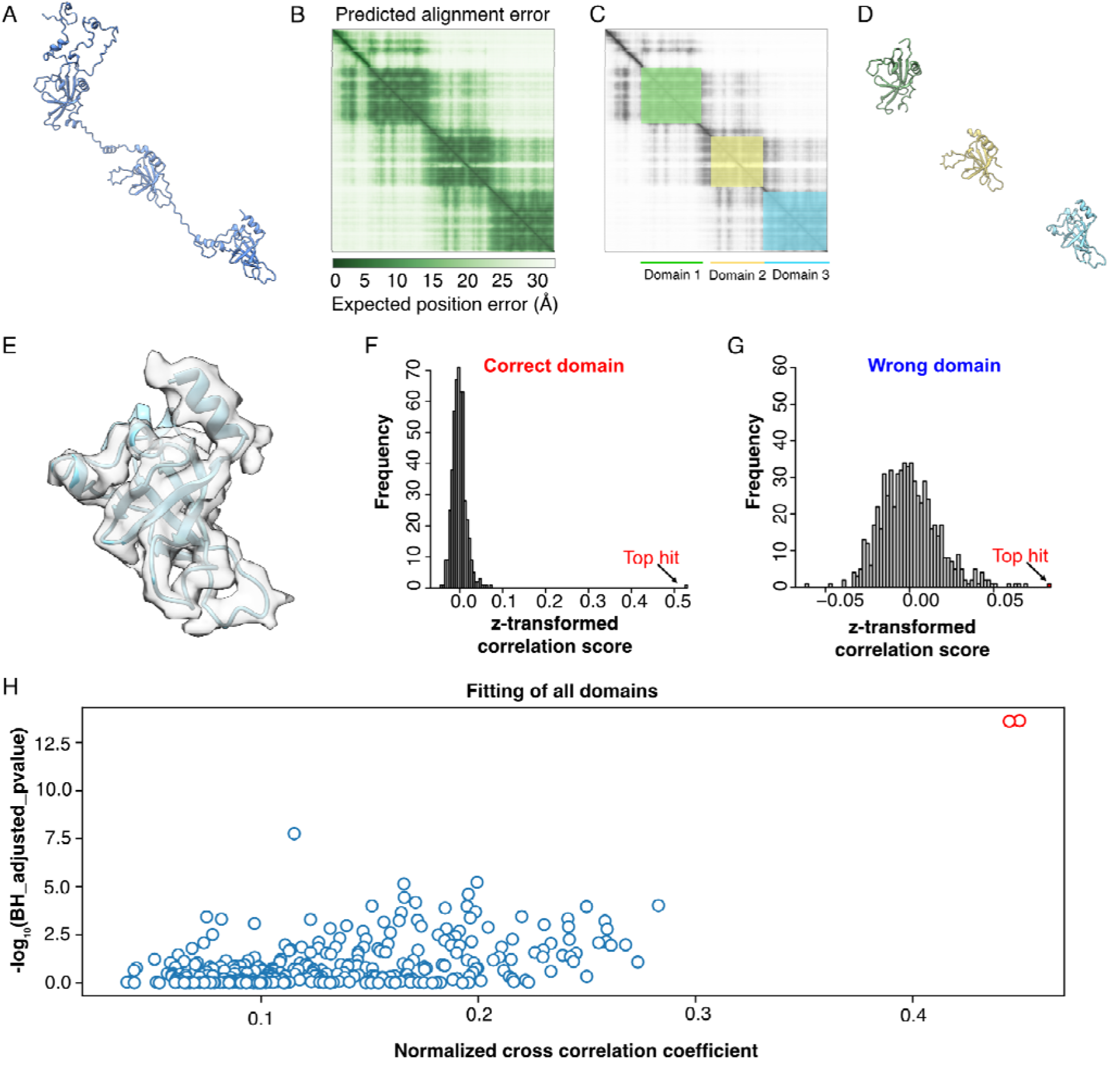
Domain partitioning and fitting evaluation in DomainFit. (A) An example of an AlphaFold2-predicted model of a multi-domain protein (RIB72B, UniprotID I7MCU1). (B) The PAE plot of the RIB72B. (C) Domain partitioning of RIB72B based on the PAE plot. (D) Domain partitioning of RIB72B corresponding to (C). (E) Fitting of a domain of RIB72B inside its corresponding segmented density at 4 Å resolution. (F) Histogram of the z-transformed correlation score of all obtained fits of the correct domain of RIB72B. The top hit (red) is separated from the score distribution. (G) Histogram of the z-transformed correlation score of a wrong domain into the same density in (F). (H) The scatter plot to visualize the value of - log_10_ of Benjamini Hochberg adjusted p-value versus the normalize cross correlation coefficient of each domain fitting into the density of interest. The top right corner two points are well separate for the rest of the point cloud, indicating likely well-matched domains into the density.

To increase the flexibility of our program, *process_predicted_models.py* writes out the information of the predicted domains using the format employed by DPAM 31. As a result, we can use domain information data from either *phenix.process_predicted_model* or existing domain data of certain organisms such as human, mouse, and zebrafish from DPAM 31 to generate the PDBs of individual domains. In the case of a small number of target proteins, we can even edit the domain information files manually to generate customized domain parsing.

At this step, we recommended imposing a filter on the minimum and maximum domain sizes for fitting. At the lower end, 40 amino acids is the minimum size that allows us to reduce the number of domains used without affecting the results since domains under 40 amino acids tend to be either wrongly partitioned or not compact. On the upper limit, we recommend using a bigger value like 1,000 amino acids as a fail-safe in the case that domain parsing does not work properly.

To prepare for the fitting of domains, a volume containing a compact density should be segmented from the cryo-EM map. The compact density means either globular or coiled-coil bundle. A long helical density does not work since its shape is not unique for identification of protein fold. Ideally, the segmented density should be equivalent to a domains of more than 50-300 amino acids in size. This serves two purposes: (i) to reduce computational time, and (ii) to avoid the wrongly predicted tertiary conformation caused by long flexible loop between different domains.

Once the segmented density is prepared, *fit_domains_in_chimerax.py* is used to fit all the individual domain models into the density to find the best fitting position of each domain (Fig. 1A, Fig. 2E). The script uses the ChimeraX *fitmap* command to fit each domain in the density from a fixed number of initial random translated and rotated search positions within the density defined as “initial search placements”. Instead of exhaustive six dimensional search like Colores package in Situs, for global search, ChimeraX *fitmap* places the model randomly in rotation and translation within the map, and performs rigid-body local optimization from each initial placement. After the local optimization, ChimeraX will group the fitting solutions of based on similarity in orientation. Therefore, the number of fitting solutions tend to be smaller than the number of initial search positions due to clustering. These fitting solutions’s correlation score are then z-transformed and then used for the calculation of the p-value, the likelihood that the top hit is correct and clearly significant compared to the rest of the fitting solutions (See Materials & Methods, Fig. 2F). Once the fitting is done for every domain in the density, we expect that top hits with the correct fold/domain are clearly distinguished from other fitting solutions by their z-transformed scores, whereas those with the incorrect domain are not so differentiated (Fig. 2G). Visualizing the fitting statistics i.e. p-values and cross correlation coefficient of all domains allow a quick way to see how distinct is the top hits (Fig. 2H).

A comma-separated value (CSV) file containing the best fitting position of each domains is written at the end, sorted first by p-value and then by the correlation score for quick evaluation.

### DomainFit benchmarking for MIP identification

To assess the efficiency of DomainFit, we tested it on segmented densities from the single-particle cryo-EM map of the doublet microtubule from *Tetrahymena thermophila* (EMD-29685) at ∼4 Å resolution. More than 40 MIPs were identified and modelled using an AI-assisted modelling approach (Fig. 3A). Even though the resolution of the doublet microtubule cryo-EM map was high, modelling and identifying proteins was laborious and time-consuming. Our goal for this test was to see whether we could identify the MIPs using DomainFit independently.

**Figure 3:**
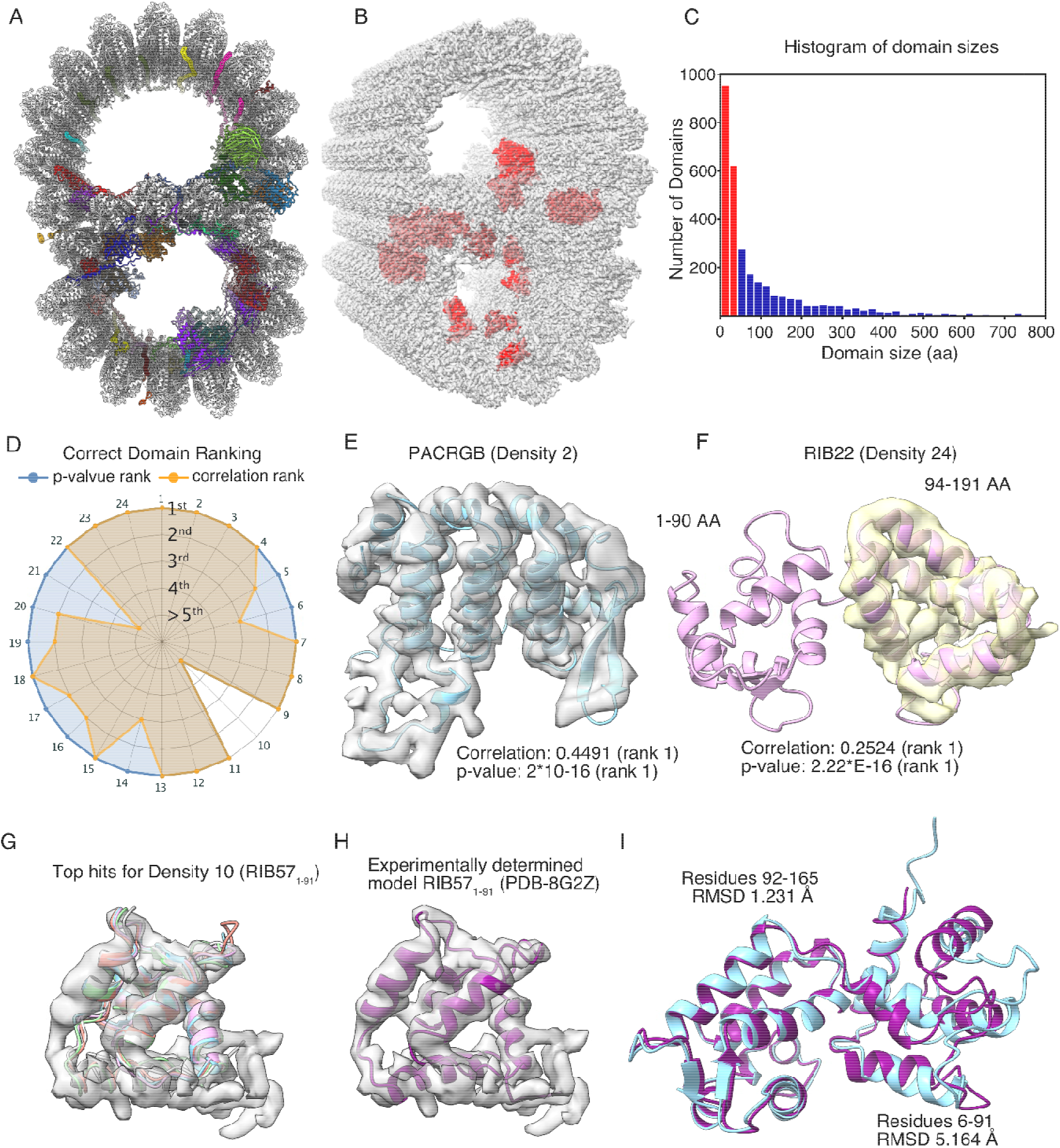
Identification of MIPs in the *T. thermophila* doublet microtubule. (A) The doublet microtubule structure with MIPs (PDB: 8g2z). (B) Densities (red) in the cryo-EM map of the doublet microtubule (EMD-29685) for testing with DomainFit. (C) A histogram of domain size. Red bars indicate domains of less than 40 amino acids and are excluded from fitting. The x-axis is limited to a domain size of 800 amino acids. (D) DomainFit ranking by p-value and correlation about mean of the correct domain for 24 densities. (E) Example of the perfect match between the AlphaFold2-predicted model of PACRGB fitted within the corresponding density. (F) The domain of RIB22 is not partitioned as expected. However, the bigger domain was still fitted correctly into the density due to good agreement between the AlphaFold2-predicted model and the density. Due to incorrect parsing and the fact that the EF-hand domain is abundant in the cilium, the fit of the correct domain is ranked 9 but the p-value is still ranked 1st. (G) When the fold of the density is common (EF-hand domain RIB57_1-91_ - Density 10), the top hits consist of domains with a similar fold. The top five hits are all domains from CFAP115, which have the same EF-hand fold as the correct domain RIB57_1-91_. (H) The experimentally determined model of RIB57 (PDB: 8g2z) is shown inside its corresponding density. (I) There is a big discrepancy between the experimentally determined model (cyan) and AlphaFold2-predicted model (purple) of RIB57_1-91_ (RMSD: 5.164 □), which leads to the poor fitting of RIB57.

We segmented 24 densities from 13 identified MIPs in the cryo-EM map corresponding to different domains of MIPs (Supplementary Table 1, Figure 2B). We established a database comprising 856 AlphaFold2-predicted models based on the mass spectrometry data of the WT cilia without membrane (994 detected proteins) 21. One hundred and thirty eight proteins were not available from the AlphaFold DB because of a lack of predictions due to their high molecular weight. For benchmarking purpose, we ignored them for this test since the 856 AlphaFold2-predicted models covered all the tested MIPs.

Using *process_predicted_models.py* and *save_domains_from_info.py*, we divided 856 AlphaFold2-predicted models into 2819 domains. We removed domains of less than 40 amino acids which left us with 1561 domains for fitting (Fig. 3C). We then fitted all 1561 domains into each density filtered at 4 Å using 200 initial search position placements. On average, each density search took ∼30 minutes using 12 cores (20 threads) on an i7-12700K (12 core-CPU and 32GB RAM).

Out of all the densities tested, we found 22 out of 24 correct domains are present in the top 5 hits ranked by descending p-value (Supplementary Table 1, Fig. 3D, Fig. S1). Overall, we found the p-value to be a better predictor of correct identification and fitting than the normalized cross correlation coefficient, which is called correlation about mean in ChimeraX. Often, the domains were correctly found as the top hits with the highest correlation and best p-value such as PACRGB, an inner junction protein 32 (Fig. 3E). Our results show that the p-value ranks consistently higher than the correlation score for the correct domains for the densities (Fig. 3D). In the cases where domains have a common fold, multiple domains returned a p-value of 2.22*10^−16^ (limit of R program). Thus, we sorted the fitting solutions first based on p-values, and then by their normalized cross correlation coefficient. As a result, hereafter, top hits will refer to the top p-value ranked hits.

In some cases such as Density 6 (CFAP52_13-317_), by looking at the fit of the domains in the density, we can roughly estimate the size of the domain and establish a minimum domain size threshold, which allows quicker identification of the correct domain (Fig. S2A-B).

There are quite common cases where the partitioned domain is bigger than the segmented density (Supplementary Table 1, Fig. S1, Density 6, 10-13, 16-17, 19-22, 24). This might affect the fitting of DomainFit. For example, our parameters for domain parsing tend to keep entire EF-hand domain pairs as a single domain, instead of two separate EF-hand domains. For RIB22, while the density represents only RIB22_1-91_, RIB22_4-191_ was partitioned as one domain (Fig. 3F). On the other hand, the correct fitting was found for RIB22_4-191_ with a p-value and correlation value of rank 1^st^ (Fig. 3F, Supplementary Table 1).

Interestingly, for Density 10, corresponding to the EF-hand domain RIB57_1-91_, DomainFit failed to find the correct domain (Fig. 3G). While all the top hits were the wrong protein, they all had the correct EF-hand fold (Fig. 3G). when we compared the experimentally determined model of RIB57 (Fig. 3H) and the AlphaFold2-predicted model, there was a big difference between the two models from residues 6 to 91 (RMSD 5.164 A, Fig. 3I). Therefore, the failure to find RIB57_1-91_ density (Density 10) was due to both the difference in the AlphaFold2-predicted model and the unsuccessful parsing of RIB57_1-91_ domain.

We conclude that DomainFit works well but attention needs to be paid to potential mis-identification of proteins due to their size and domain parsing. Also, running with a higher number of initial search placements is a safeguard against inaccuracy in domain parsing to accommodate more initial translated and rotated positions.

### Performance of DomainFit depending on the number of initial placements and resolution

Our test results showed that DomainFit is effective in identifying compact densities and the p-value is the most reliable indicator of the correct hit. For p-value to be estimated properly, a reasonable number of data points, i.e. fitting solutions are needed. Therefore, we wanted to explore the sensitivity of the p-value estimation to the number of initial search placements. We ran DomainFit for Density 1 100 times for each initial search placement value ranging from 1 to 300 and plotted the average p-value found. Our analysis shows that above 150 initial search placements, the resulting p-value is consistently found to be as high a value as it can be (Fig. 4A). Thus, setting a search placement value of 150 should yield optimal results for the speed and accuracy of the fits in this setup. This setting is appropriate for a domain of 100-150 amino acids while larger domains may require a higher number of initial search placements. We also looked at the z-transform correlation score for the correct domain at different numbers of initial search placements (Fig. S3A-D). The statistical calculation is certainly less reliable when the number of fitting points is small. For the correct domain, if the correlation score of the top hit is distinct from that of the rest, a good p-value can still be obtained.

**Figure 4:**
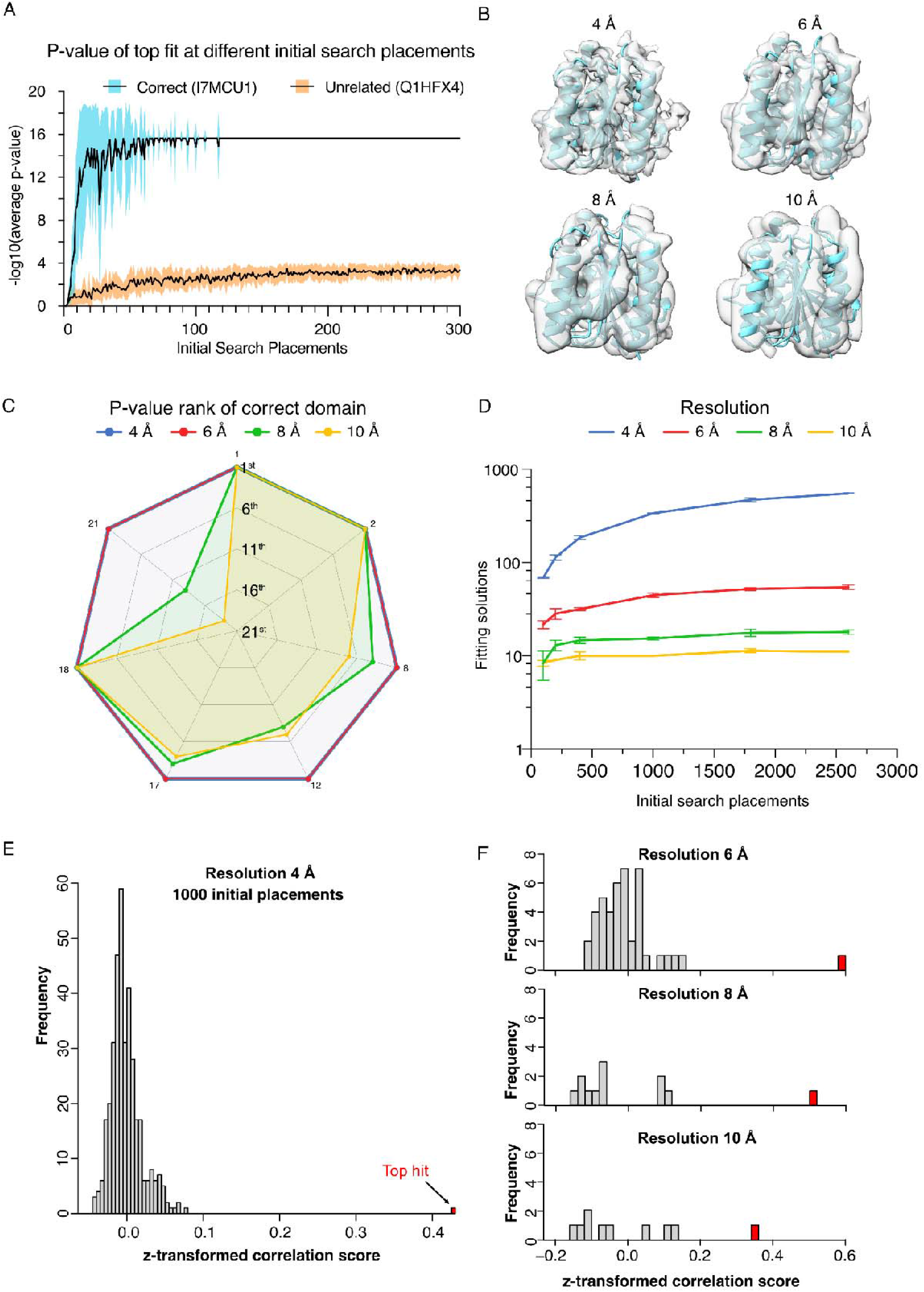
Performance of DomainFit at different numbers of initial search placements and resolutions. (A) -log10 of the average p-value at increasing initial search placements for a correct and an unrelated domain. Standard deviations are shown in coloured regions. (B) Appearance changes of a density filtering at 4, 6, 8 and 10 Å resolution. (C) p-value rank of the correct domains fitted to different densities of MIPs at 4, 6, 8, and 10 Å resolution. (D) Number of fitting solutions versus number of initial search placements for maps at 4 Å (blue), 6 Å (red), 8 Å (green) and 10 Å (yellow) resolution. (E) A histogram of the z-transformed correlation score of a correct domain fitted to a density at 4 Å resolution shows a clear separation between the top hit and the rest of the fitting solutions. (F) Histograms of z-transformed correlation score of the same domain fitted to the densities filtered at 6, 8 and 10 Å resolutions. There are significantly fewer fitting solutions at lower resolutions and the separation of the top hit, and the rest is reduced.

The number of fitting solutions found does not depend only on the number of initial search placements but also on the resolution of the map. Next, we wanted to check the effect of map resolutions on the success of DomainFit. To test whether the program could successfully identify domains at lower resolutions, we filtered and ran the density maps of the MIPs at 4, 6, 8, and 10 □ (Fig. 4B). We found that the search consistently identified the correct domain within the top 10 results at 4 to 8 □ resolution (Fig. 4C). At lower resolutions, there are significantly fewer fitting solutions found even with the same number of initial search placements (Fig. 4D). This is because of the smoothening of the map at lower resolutions, which does not allow many unique fitting solutions. Because of this, ChimeraX produces a small number of clusters of fitting solutions from a large number of initial search placements. This leads to less confidence in statistical analysis of the top hits. If we look at the z-transformed correlation score at 4, 6, 8 and 10 Å of the correct domain (Fig. 4E, F), we can see that at 6 Å, the distribution of the fitting solution and the top hits still maintains its shape, allowing reliable p-value calculation. At 8 Å, while the top hit is still well separated from the distribution, the rest of the solution does not look normally distributed. At 10 Å, not only does the distribution of the correlation score is not normally distributed but the difference between the top hit and the rest gets smaller.

As a result, DomainFit seems to work best in the 4-to-6 Å resolution range. At 8 Å resolution, the number of initial search placements must be increased to help with statistical calculation. At lower than 8 Å, DomainFit should only be used for finding types of folds that fit well in the density by visual inspection of the top hits but it should not be used for domain identification.

### New MIP identified in the doublet microtubule

With our success in testing and validating the DomainFit program, we attempted to identify unknown proteins from cryo-EM maps with a resolution lower than 4 Å. An unidentified MIP was previously reported in the subtomogram average of the *T. thermophila* doublet microtubule 33 (Fig. 5A). However, this density was not observed in the single particle cryo-EM map of the doublet microtubule 21 (Fig. 5B) and at too low resolution to be identified.

**Figure 5:**
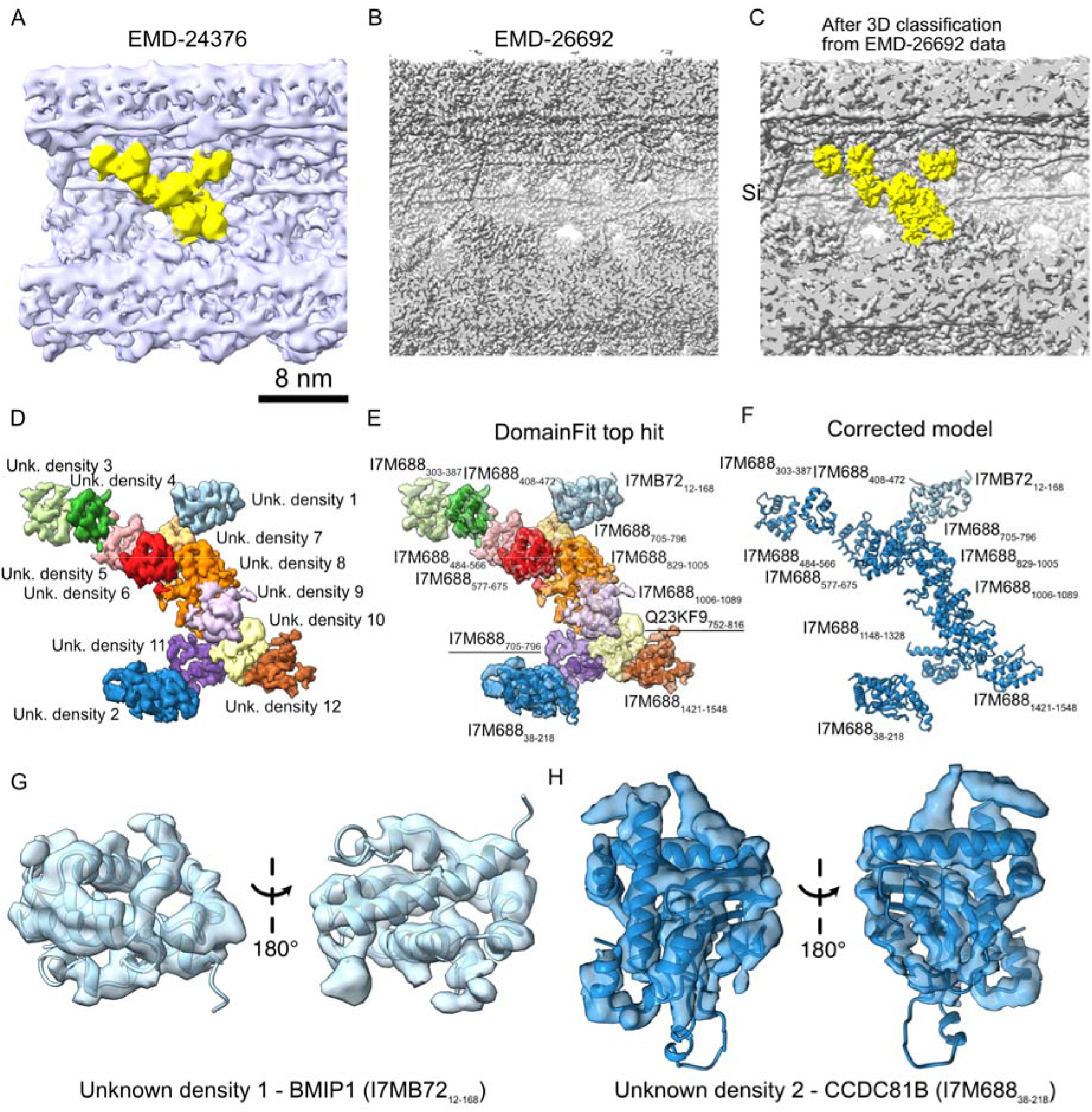
New proteins identified from the doublet microtubule of *T. thermophila.* (A) Overview of the unknown density (yellow) from the 48-nm repeating unit subtomogram average map of doublet microtubule (EMD-24376) 33. (B) The same density is not visible in the same view of the single particle cryo-EM map of the 48-nm repeating unit of the doublet microtubule from the *T. thermophila* K40R map (EMD-29692) 21. (C) The unknown density exists in the doublet microtubule map after 3D classification. (D) Segmentation of the unknown density into different small unknown densities for DomainFit. (E) DomainFit top hits for each unknown density using an AlphaFold database of salt-treated ciliome. (F) Corrected model of protein domains after examining, showing BMIP1 (UniprotID I7MB72) is density 1 and CCDC81B (UniprotID I7M688) forms density 2-12. (G) Fit of AlphaFold2-predicted model BMIP1_12-168_ into density 1. (H) Fit of AlphaFold2-predicted model CCDC81B_38-218_ into density 2.

We re-processed the cryo-EM dataset containing the 48-nm particles of *T. thermophila* K40R doublet microtubule 21 with Cryosparc 34 Using 3D classification and refinement, we obtained a 4.5 Å resolution cryo-EM map with the same density as the subtomogram average (Fig. 5C, Fig. S4A, B).

To identify the proteins in the unknown density, we segmented the density into 12 smaller densities (Fig. 5D). To improve the accuracy of the search, we used the ciliome of the salt-treated doublet microtubules 4. The reason is that the cryo-EM map of salt-treated doublet microtubules retains most of the MIPs and the unknown density 35. The salt-treated ciliome of 166 proteins was divided into 334 domains of at least 40 amino acids. We ran DomainFit with these 334 domain models for the above 12 densities.

Interestingly, the top hits for the 12 densities came from only three proteins (Fig. 5E, Supplementary Table 2). Upon inspecting the results, we concluded that the 12 densities are composed of only two proteins I7MB72 (TTHERM_00525130) and I7M688 (TTHERM_00649260) because of the overall architecture (Fig. 5F, Fig. S5A) and the fit of unique domains in both proteins (Fig. 5G-H).

I7MB72 matches unknown density 1 with high confidence in p-value and correlation (Fig. 5F), referring as BMIP1 from here onwards. Upon inspection of the map, density 1 was always present in the doublet microtubule (EMD-29692) but was not identified previously. As a result, we could model extra regions of the protein BMIP1 in the *T. thermophila* K40R map (EMD-29692) at 3.5 Å resolution, confirming the identity of unknown density 1 and the success of DomainFit (Fig. 6A, B). In addition, BMIP1 is abundant in both salt-treated and native doublet microtubules similar to the profiles of most MIPs. BMIP1 has a cross-link to CFAP45, which exists as two copies between protofilaments B8B9 and B7B8, as shown in studies that usedchemical cross-linking coupled with mass spectrometry of cilia 21,36 (Fig. 6C). While the cross-linked residues (Lysines 213 and 222) from BMIP1 are not modelled, they seem to be within the cross-link distance (< 35 Å) from lysines 403 and 283 of CFAP45.

**Figure 6:**
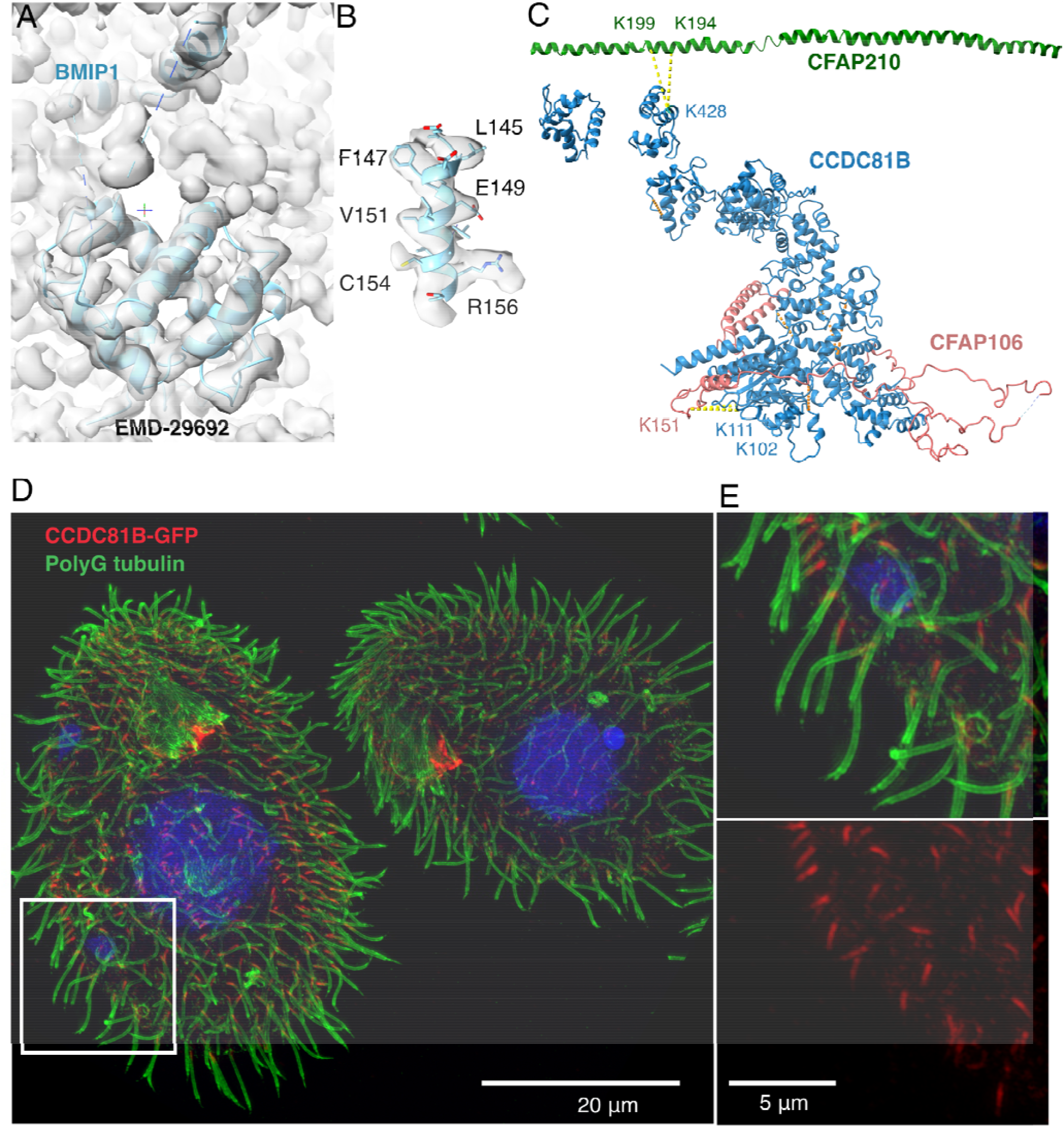
Validation of the newly identified MIPs. A. Model of BMIP1 (UniprotID I7MB72) using Coot. B. Side-chain densities of a helix from BMIP1. C. Intra-cross-links (yellow) within CCDC81B and inter-cross-links (orange) between CCDC81B and CFAP106, a known MIP protein 32. D. Merged super resolution-structured illumination microscopy image of *T. thermophila* cells with CCDC81B-GFP shows that CCDC81B only localizes to the proximal region of the cilia. Rectangle indicates the zoom-in view in (E). Red: CCDC81B-GFP signal; Green: doublet microtubule labelled by polyG-Antibody; Blue: Hoechst dye labelling the nucleus. E. The zoom-in merged image (top) and GFP channel (bottom) showing localization of CCDC81B to the proximal region.

Densities 2-12 (Fig. 5D) belong to I7M688, a CCDC81-domain containing protein. We named I7M688 as CCDC81B from here onwards. The CCDC81 domain of CCDC81B is unambiguously fitted into unknown density 2 (Fig. 5H). There are a few CCDC81-containing proteins in the cilia of *T. thermophila*: IJ34 (UniprotID I7M9T0), CCDC81A (UniprotID I7MLF6), CCDC81B (UniprotID I7M688), Q240Y1 and Q22HG4. IJ34 is a MIP near the inner junction 21 and CCDC81A was identified as a ciliary tip protein 37. Q240Y1 and Q22HG4 are clear paralogs of CCDC81B with similar architecture but are washed away in salt-treated cilia 21.

Mass spectrometry results show that CCDC81B is resistant to salt and is similar to a MIP profile 21. *In situ* cross-linking mass spectrometry shows that CCDC81B is crosslinked to tubulin alpha, CFAP106, FAP210, and interestingly RIB43A 36. All these cross-links are satisfied relative to the CCDC81B position in the unknown density, except for RIB43A (Fig. 6C). Therefore, we are confident that CCDC81B corresponds to unknown densities 2-12.

For a quick comparison of DomainFit with complete AlphaFold2 model fitting, we also fitted 166 complete AlphaFold2-predicted models into the full segmented density (Fig. S4). The top hits were IJ34 and CCDC81B. Both proteins contain a CCDC81 domain (Fig. S5B). BMIP1 only ranks 22 in p-value. The AlphaFold2-predicted model of CCDC81B has good tertiary structure prediction, which helps the complete model fitting. However, this shows the weakness of complete model fitting. First, complete model fitting seems to penalize smaller domains such as BMIP1. Second, doing complete model fitting does not allow to filter the sizes of the fitted domains, which is useful to eliminate false positive fits. Finally, with DomainFit, we can visualize the top hits from different densities (Fig. S5A), allowing a quicker identification of multidomain proteins without relying on the quality of the tertiary structure prediction. On the other hand, complete model fitting of 166 AlphaFold2-predicted models in the density filtered at 10 Å resolution also found CCDC81B as the top hit with good fitting accuracy, thanks to good tertiary structure prediction (Fig. S5C, D).

To further confirm that CCDC81B is a MIP found only in a subset of the doublet microtubule, we generated a *T. thermophila* strain with CCDC81B fused with GFP. Under superresolution structured illumination microscopy, the signal of CCDC81B was limited to the 1-1.5 um proximal region of the cilia (Fig. 6C). That explains the ratio of about substoichiometric fraction of the particles containing CCDC81B densities from the single-particle cryo-EM data. As a result, we can confirm that CCDC81B is a MIP that localizes to the proximal region of the cilia.

## Discussion

In this paper, we demonstrated that it is possible to identify compact domains of proteins in maps at a resolution lower than 4 Å using the DomainFit program. We benchmarked DomainFit using known structures and successfully identified CCDC81B and BMIP1 as two new proteins in the doublet microtubule of *T. thermophila*. Notably, we showed that CCDC81B only exists in the proximal region of the cilium using superresolution structured illumination microscopy of *T. thermophila* cells expressing CCDC81B fused to GFP.

While DomainFit works, it is not as definite without complementary data as high-resolution cryo-EM data where side-chain information is available. We showed that the statistical analysis of the best-fitting domain works well. When the fold of the domain is unique, the identification works excellently. When the fold is not unique, DomainFit still finds the correct fold. Tools like Foldseek 38 can then be used to list all the proteins with the same fold available. With other complementary info such as chemical cross-linking mass spectrometry, BioID, and stoichiometry of proteins from quantitative mass spectrometry, it is possible to narrow down and identify the right proteins as demonstrated by this work and other integrated structural biology approach 39. There are certain cases in which the AlphaFold2-predicted model is not similar to the protein fold in the map, leading to the wrong identification. However, it does not happen often with compact domains.

The validation of DomainFit using known MIPs highlights a caveat of the workflow in the partitioning of domains. Breaking proteins into compact domains will never be perfect since over-partitioning will produce smaller domains and therefore increase false positive fits while under-partitioning produces bigger domains, requiring a high number of initial search positions, compromising the computational requirement. Perhaps, the development of domain partitioning with better customization in the future can improve the usability of DomainFit. Despite that caveat, our work shows that domain fitting is in general more accurate than complete model fitting since AlphaFold2 domain prediction is in general more accurate than AlphaFold2 tertiary structure prediction.

Another point to consider for a successful identification is the quality of the map. It seems that DomainFit works well between 4 and 6 Å resolution where detailed secondary structural features can be visualized. It is recommended to use appropriate post-processing methods to improve the interpretability of the map such as DeepEmhancer 40, LocSpiral 41 or Density Modification 42.

At the lower resolution of 8-10 Å, DomainFit can still work but requires complementary data for validation. For lower resolution, perhaps using domains with bigger partitions or even the entire AlphaFold2-predicted model might help with the fitting and identification if the tertiary structure is predicted correctly. With the flexibility of DomainFit, users can try both approaches and evaluate the fitting visually in the case of low resolutions using scripts provided by DomainFit.

In addition, since the program is rather quick, it is possible to use DomainFit to construct the structures of complexes with known compositions. A database of domains from all the known components can fit into different densities of the map. This allows a more unbiased way of building up the structure of the protein complex.

In conclusion, we presented here the DomainFit program, which allows the unbiased fitting of domains of proteins into the cryo-EM map. At resolutions better than 6 Å, the program can be used reliably as a tool to identify compact proteins in the map.

## Materials & Methods

### Domain parsing

For domain parsing, we used the option *split_model_by_compact_regions=True* from *phenix.process_predicted_model*. In addition, we set the option maximum_domains for each protein so that the maximum_domains equal the protein size in amino acids divided by 100 amino acids.

### p-value calculation

The p-value calculation for the fitting of each domain has been implemented in R 43 in the integrated modelling software Assembline 45 and used for the identification of the nuclear pore complex Y-shaped scaffold 44.

In brief, for each domain, ChimeraX clusters the results into a number of fitting solutions with distinct rotational and translational parameters, and cross-correlation scores. As a result, many searches with different initial rotation and translation placement, resulting in similar final fitting orientation are clustered into one fitting solution. The correlation scores of all fitting solutions are transformed to z-scores using Fisher’s z-transform and centered to yield an approximately normal distribution (Fig. 2F).

Then, the two-sided p-value and Benjamini-Hochberg adjusted p-value are calculated from the z-scores using the false discovery rate (fdr) package in R. The top hit for each domain which corresponds to the highest correlation score and lowest p-value is recorded in an aggregated list. Once this process is done for all the domains, the list is then sorted by p-value. When the correct domain is evaluated, the top hit’s z-transformed score is clearly discriminated from the rest of the fitting solution while the difference in value is not as clear when a wrong domain is evaluated (Fig. 2F, G). That translates to a lower p-value for the top hit of the correct domain.

The p-value is a more reliable indication of true positives than the correlation score since it is less sensitive to the size of the domain fitted in the density. Small domain models due to wrong partitioning always have high correlation scores because they fit perfectly in a small area of the density. For the correct domain, the correlation score difference between the top hit and the second hit tends to be larger except for domains with a symmetrical shape (e.g. WD40 beta-propeller). We also found that the Benjamini-Hochberg adjusted p-value functions similar to p-value.

### Cryo-EM and image analysis

The single particle cryo-EM data of the 48-nm repeating unit of the doublet microtubule from *T. thermophila* was originally published in 21.

To obtain the density with a resolution better than 12 Å resolution of the subtomogram average, we constructed a focused classification mask at the region of the density based on the EMD-24376 subtomogram average map and performed 3D classification without alignment for four classes on the 48-nm particles cryo-EM dataset of *T. thermophila* K40R doublet microtubule 21 using Cryosparc 34 (Fig. S4A). After classification, class 1 contains ∼45,000 particles showing the same density as the subtomogram average (Fig. 5C). This suggests that the density feature observed by subtomogram averaging is not uniformly located in the cilia. We further refined Class 1 in Cryosparc with focused local refinement using a refinement mask slightly larger than the classification mask, resulting in a cryo-EM map of 4.5 Å resolution (Fig. 5C, Fig. S4A, B). The details in the density map suggest a 5-5.5 Å resolution.

The final map was post-processed using DeepEMhancer 40.

All the visualization of maps and models were done in ChimeraX 46.

### Cryo-EM density map segmentation

To segment the density of interest for DomainFit from the cryo-EM map, we manually placed markers onto the density of interest in ChimeraX. After that, we colored the density around the markers, and use *volume splitbyzone* function of ChimeraX to segment the density out and save.

### Modelling

For modelling of BMIP1 (UniprotID I7MB72, TTHERM_00525130), we started with the AlphaFold2-predicted model of the globular domain of BMIP1 fitted in its density. We fixed the model manually in Coot 47 and modelled some extra regions of the protein. The final models were then real-space refined in Phenix 46.

For the modelling of CCDC81B (UniprotID I7M688, TTHERM_00649260), best fitting positions of different domains of CCDC81B were found using DomainFit. We joined all the domains together using Coot 47 and refined in Phenix 36.

### Cross-link mass spectrometry visualization

Cross-links to BMIP1 and CCDC81B were obtained from a chemical cross-linking mass spectrometry of *T. thermophila* cilia (Reported in Supplementary Table 1 in 48). The crosslinks were visualized in ChimeraX using the bundle XMAS 49.

### Cell culture and gene editing

All *Tetrahymena* strains used in this study were grown in SPP media 50 in a shaker incubator at 30°C and 120 rpm.

The CCDC81B gene, *TTHERM_00649260* gene was edited by homologous DNA recombination using a targeting plasmid carrying the neo4 selectable marker. The portions of *TTHERM_00649260* required for gene targeting were amplified using primers 5F (5’- ctatagggcgaattggagctttgtgaaatagatggaagag-3’) and 5R (5’- atcaagcttgccatccgcggacttgtgaatttttaaagagat-3’) amplified a terminal portion of the coding region and primers 3F (5’- gcttatcgataccgtcgaccatcaattatttcaaagtattaa-3’) and 3R (5’- agggaacaaaagctgggtacgcattatccaaaatatattctaa −3’) amplified a portion of the 3’ UTR and cloned into the pNeo24-GFP plasmid 51. The resulting edited fragment *TTHERM_00649260* was targeted to the native locus using biolistic bombardment of *T. thermophila* cells and paromomycin selection.

### Immunofluorescence

For immunofluorescence, *T. thermophila* cells were fixed and stained as described 52. The primary antibodies used were the mouse monoclonal anti-polyglycylated tubulin AXO49 (diluted 1:200) 52 and polyclonal anti-GFP antibodies (Rockland, 1:800). The secondary antibodies used were goat-anti-mouse IgG-FITC and goat-anti-rabbit-Cy3 antibodies (Jackson Immunoresearch). SR-SIM imaging was conducted on an ELYRA S1 microscope equipped with a 63× NA 1.4 Oil Plan-Apochromat DIC objective. The optical slices were analyzed by Fiji/ImageJ (Z project tool).

## Data Availability

The data generated in this study are available in the following databases: XXX (The proximal proteins from the 48-nm Tetra K40R doublet), <u>EMD-</u>YYY (The proximal density from the 48-nm Tetra K40R doublet). All data used but not produced in this study are available in the following databases: <u>8G2Z</u> (PDB-model of the 48-nm Tetra WT doublet), <u>EMD-29685</u> (48-nm Tetra WT doublet), <u>EMD-29692</u> (48-nm Tetra K40R doublet).

## Code Availability

The code for DomainFit is available at https://github.com/builab/DomainFit

## Acknowledgements

We thank Drs. Mike Strauss and Corbin Black for critically reading of the manuscript. KHB is supported by grants from the Canadian Institutes of Health Research (PJT-496210) and the Natural Sciences and Engineering Research Council of Canada (RGPIN-2022-04774). JG is supported by NIH grant R01GM135444.

## Author Contributions

Conceptualisation: KHB. Methodology: MT, JGao, TL, KHB. Software: MT, JGao, KHB. Formal analysis: JGao, MT, TL, KHB. Investigation: JGao, MT, TL, CL, JG, KHB. Resources: JG, KHB. Supervision: TL, JG, KHB. Writing–original draft: TL, KHB. Writing–review & editing: JGao, TL, JG, KHB.

## Supplementary Figures

**Fig. S1:**
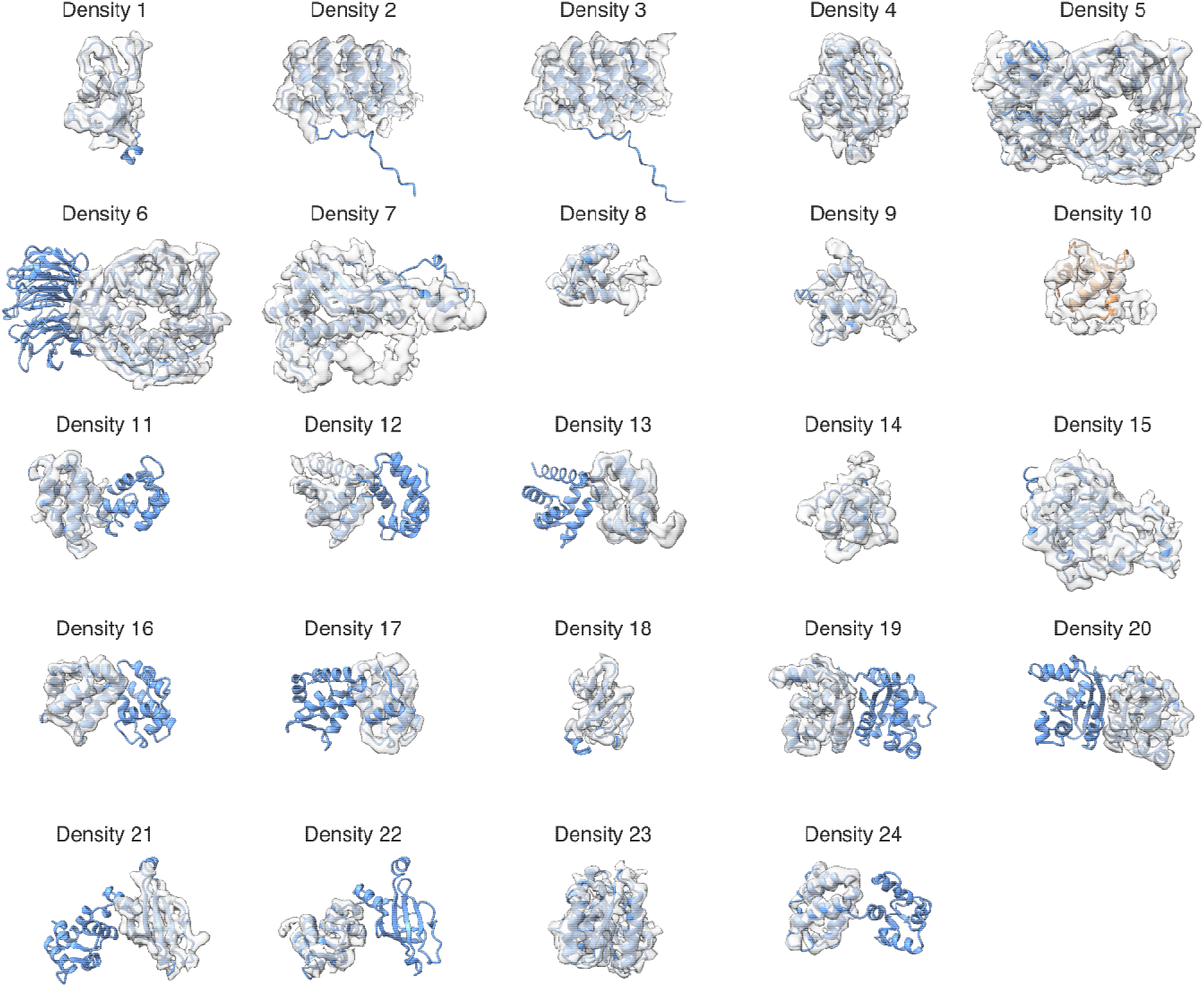
All segmented densities and best-fitted domains found by DomainFit. Blue: correct domain; Orange: incorrect domain. The correct domains corresponding to Density 6, 10-13, 16-17, 19-22, 24) were partitioned bigger than the densities.

**Fig. S2.**
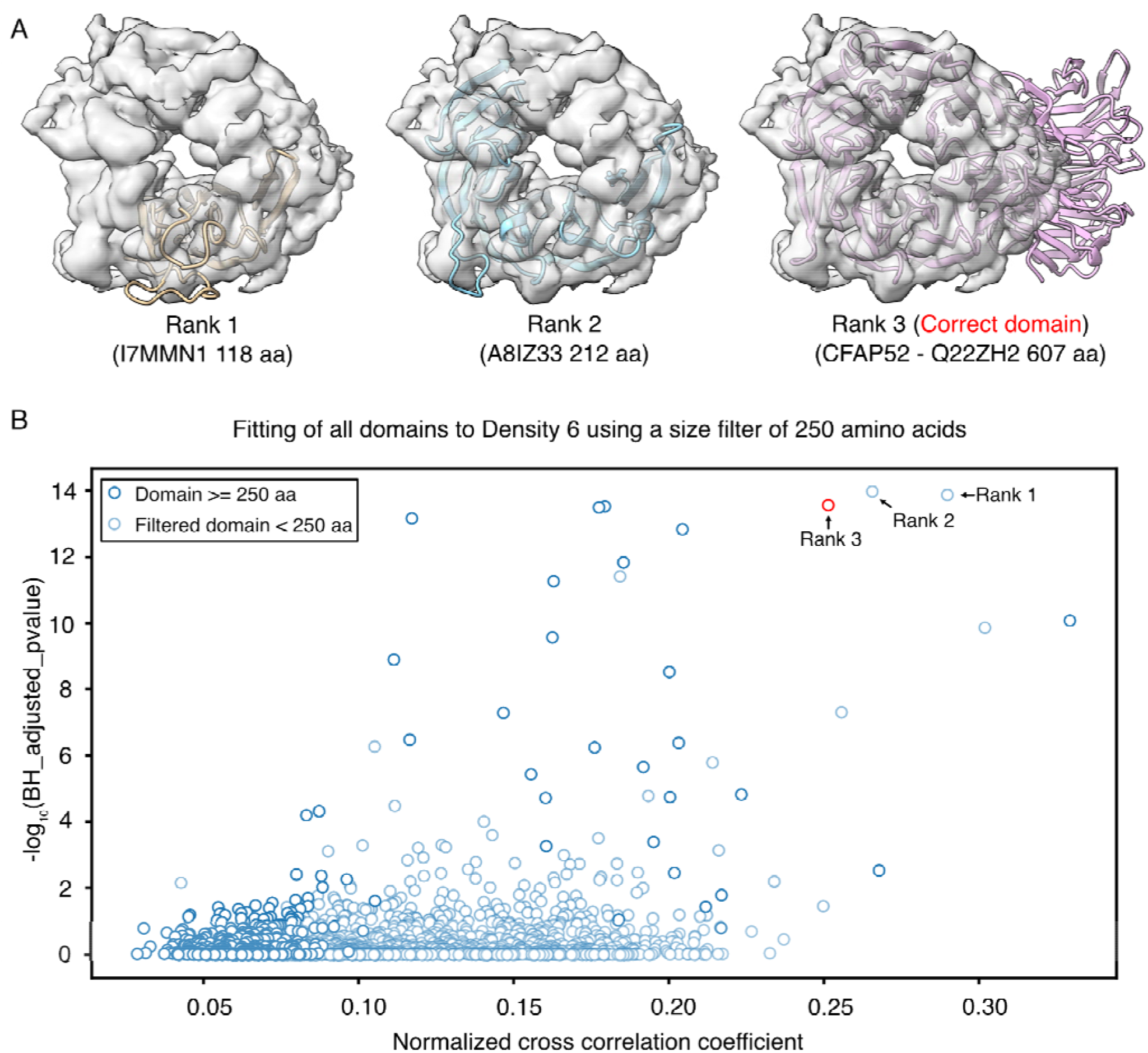
Application of domain size filtering for domain identification. (A) Top three hits for Density 6 fitting. While rank 1 and 2 domains fit well in Density 6, they are clearly not the right domain due to the unoccupied region. With the size of rank 2 domain (212 aa), it is possible to guess that domain occupied Density 6 should be at least 250 amino acids. (B) Plotting of the fitting statistics for all domains in Density 6 using a size filter of 250 amino acids. Purple indicates domain larger or equal 250 amino acids while light cyan indicates domains smaller than 250 amino acids and filtered out. With this size filtering, we can reduce false positive hits and identify the right domain easier. In this case, the correct domain (CFAP52) was ranked 3 in p-value before size filtering and ranked 1 in p-value after size filtering of 250 amino acids.

**Fig. S3.**
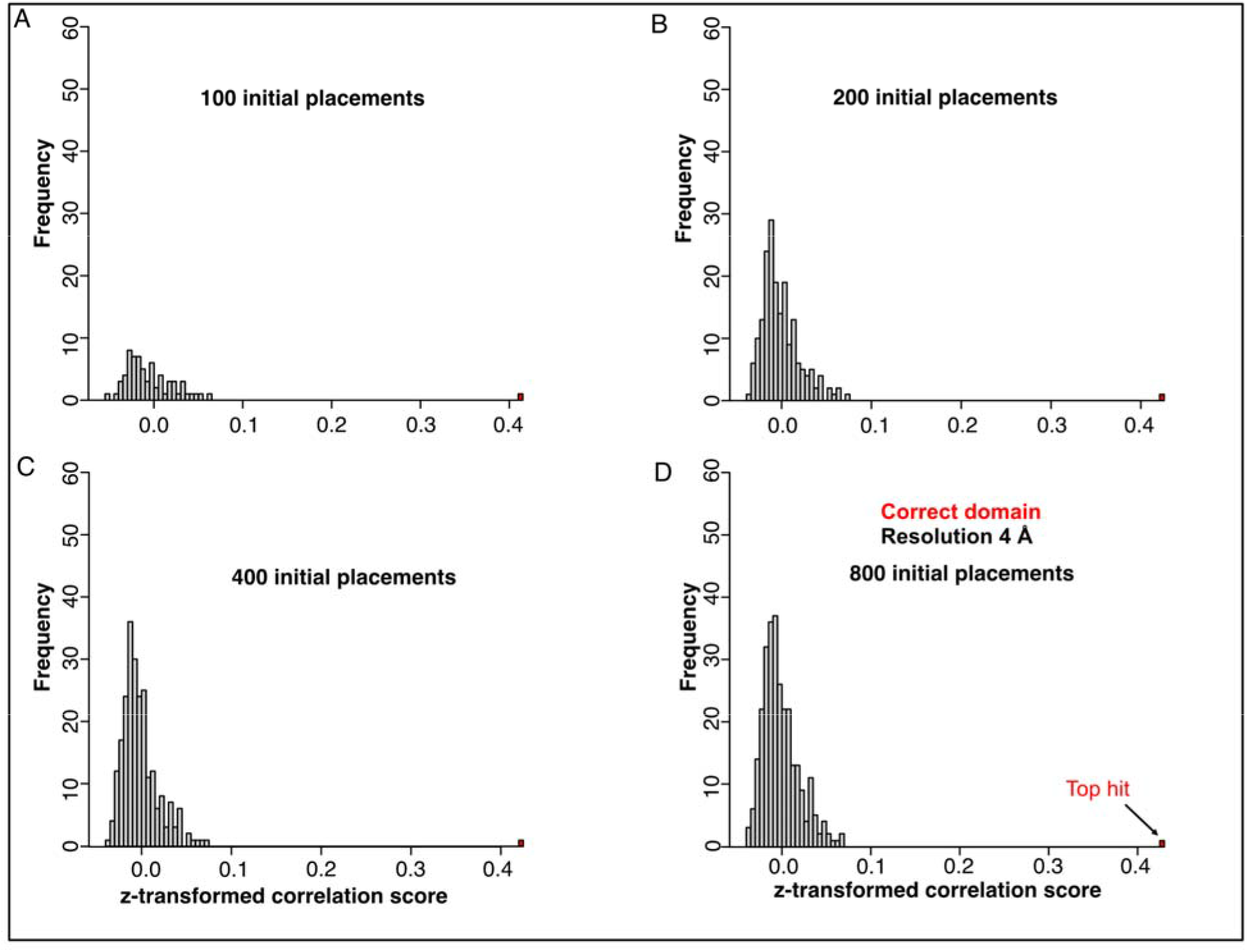
Statistical analysis of fitting solutions. (A-D) The effect of 100 (A), 200 (B), 400 (C) and 800 (D) initial search placements on z-transformed correlation score for the correct domain. Even at a sampling of 100 initial search placements, the top hit (correct solution) still seems discriminated from the rest of the fits.

**Fig. S4.**
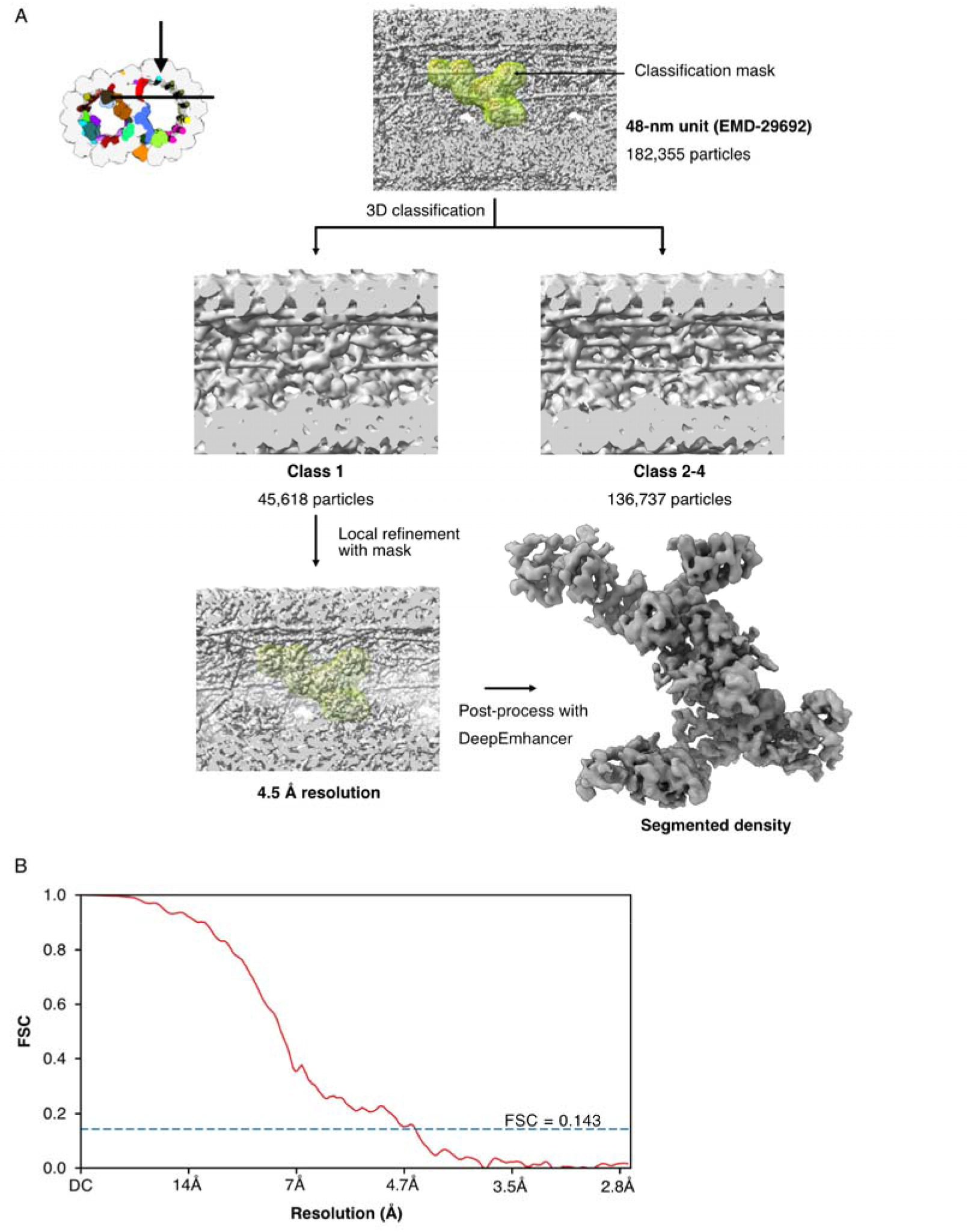
Analysis and reconstruction of the unknown density in the 48-nm doublet microtubule structure. (A) The cartoon indicates the view of the section of the B-tubule of the doublet microtubule. The classification mask is shown inside the 48-nm unit of the doublet microtubule (EMD-29692) showing the density inside is barely visible. Using Cryosparc 3D classification into four classes, Class 1 shows the density clearly inside the classification mask compared to classes 2-4. Local refinement of class 1 with the classification mask resulted in a 4.5 Å resolution structure. After post-processing with DeepEmhancer, the density is segmented out of the map for protein identification by DomainFit. (B) The Fourier Shell Correlation curve of the map obtained in (A).

**Fig. S5.**
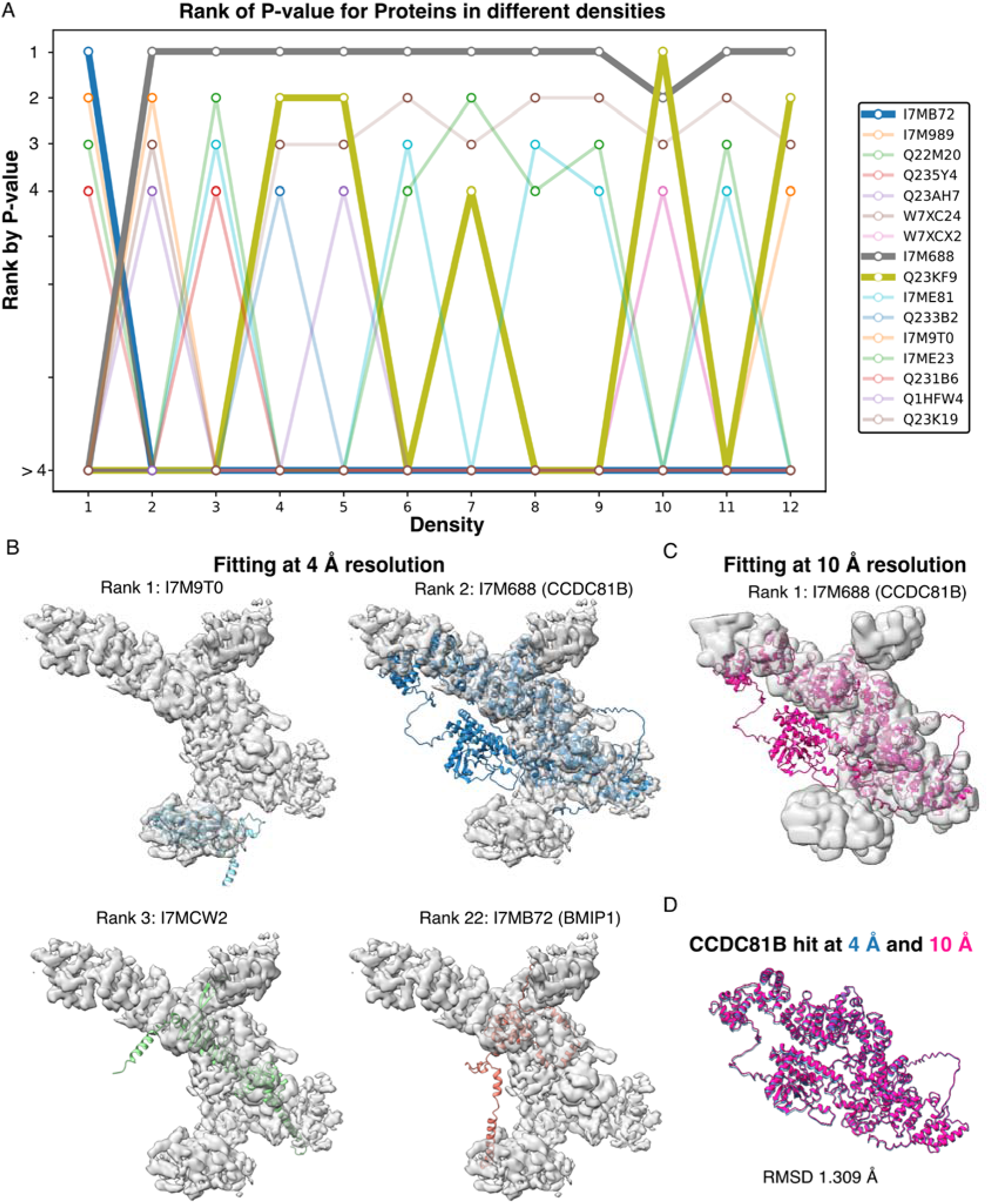
Comparison between domain fitting and complete model fitting. (A) The bump chart shows the top 5-hit proteins across 12 densities by p-value with a size filtering of 60 amino acids. Any domain ranked outside the top 4 hits is plotted at the same position in the plot (> 4). There are only three proteins among the top hits (BMIP1, CCDC81B and Q23KF9). CCDC81B is consistently the top hit in many densities, helping us to rationalize the identity of the multidomain CCDC81B. (B) Top three hits for complete model fitting into the density and the right protein BMIP1 (ranked 22). Top 1 and 2 fit well to the density while I7MCW2 and BMIP1 fit poorly in the density. (C) Complete model fitting into the proximal density filtere at 10 Å resolution. At this resolution, CCDC81B became the top hit. (D) Overlap of the best fit of CCDC81B into the proximal density at 4 and 10 Å resolution shows that the fitting positions found are almost identical (RMSD = 1.309 Å).

## Supplementary Table

**Supplementary Table 1:** Identification of MIPs in the cryo-EM map of the doublet microtubule using DomainFit.

| Density | Protein | UniProtID | Domain Residues | Corr Rank | p-value Rank | Note |
| --- | --- | --- | --- | --- | --- | --- |
| Density 1 | RIB72B | I7MCU1 | 413-516 | 1 | 1 |  |
| Density 2 | PACRGB | I7M317 | 132-319 | 1 | 1 |  |
| Density 3 | PACRGA | I7MLV6 | 1-319 | 1 | 1 |  |
| Density 4 | CFAP20 | Q22NU3 | 1-195 | 1 | 1 |  |
| Density 5 | CFAP52 | Q22ZH2 | 1-609 | 2 | 1 |  |
| Density 6 | CFAP52 | Q22ZH2 | 12-317 | 3 | 1 | Domain partitioned bigger |
| Density 7 | IJ34 | I7M9T0 | 1-251 | 1 | 1 |  |
| Density 8 | CFAP115 | Q23KF9 | 26-117 | 1 | 1 |  |
| Density 9 | CFAP115 | Q23KF9 | 133-233 | 1 | 1 |  |
| Density 10 | RIB57 | I7ME23 | 1-91 | 36 | 36 | Domain partitioned bigger, AlphaFold2 prediction different |
| Density 11 | RIB57 | I7ME23 | 92-196 | 1 | 1 | Domain partitioned bigger |
| Density 12 | RIB35 | I7ME81 | 1-96 | 1 | 1 | Domain partitioned bigger |
| Density 13 | RIB35 | I7ME81 | 99-196 | 1 | 1 | Domain partitioned bigger |
| Density 14 | RIB35 | I7ME81 | 207-295 | 3 | 1 |  |
| Density 15 | CFAP161A | Q22WJ6 | 58-280 | 1 | 1 |  |
| Density 16 | CFAP161A | Q22WJ6 | 304-389 | 2 | 1 | Domain partitioned bigger |
| Density 17 | CFAP161A | Q22WJ6 | 392-490 | 2 | 1 | Domain partitioned bigger |
| Density 18 | CFAP67 | W7XGD1 | 1-77 | 1 | 1 |  |
| Density 19 | CFAP67 | W7XGD1 | 89-221 | 2 | 1 | Domain partitioned bigger |
| Density 20 | CFAP67 | W7XGD1 | 236-371 | 2 | 1 | Domain partitioned bigger |
| Density 21 | RIB72A | I7M0S7 | 420-510 | 7 | 1 | Domain partitioned bigger |
| Density 22 | RIB72A | I7M0S7 | 526-611 | 1 | 1 | Domain partitioned bigger |
| Density 23 | RIB26 | Q232I6 | 1-237 | 1 | 1 |  |
| Density 24 | RIB22 | I7LT67 | 94-191 | 1 | 1 | Domain partitioned bigger |

**Supplementary Table 2:** Reason for identification of unknown densities.

|  | Top Hit |  | Correct domain |  | Fitting stats of correct domain |  |  |
| --- | --- | --- | --- | --- | --- | --- | --- |
| Density | UniProtID | Domain | UniprotID | Domain | Corr. rank | p-value rank | Rationale |
| Unknown density 1 | I7MB72 | 12-168 | I7MB72 | 12-168 | 1 | 1 | Unique domain |
| Unknown density 2 | I7M688 | 38-218 | I7M688 | 38-218 | 1 | 1 | Unique domain |
| Unknown density 3 | I7M688 | 303-387 | I7M688 | 303-387 | 1 | 1 | Good fit & continue with other domains |
| Unknown density 4 | I7M688 | 408-472 | I7M688 | 408-472 | 1 | 1 | Good fit & continue with other domains |
| Unknown density 5 | I7M688 | 484-566 | I7M688 | 484-566 | 1 | 1 | Good fit & continue with other domains |
| Unknown density 6 | I7M688 | 577-675 | I7M688 | 577-675 | 1 | 1 | Good fit & continue with other domains |
| Unknown density 7 | I7M688 | 705-796 | I7M688 | 705-796 | 1 | 1 | Good fit & continue with other domains |
| Unknown density 8 | I7M688 | 829-1005 | I7M688 | 829-1005 | 1 | 1 | Good fit & continue with other domains |
| Unknown density 9 | I7M688 | 1006-1089 | I7M688 | 1006-1089 | 1 | 1 | Good fit & continue with other domains |
| Unknown density 10 | Q23KF9 | 752-816 | I7M688 | 1148-1328 | 19 | 1 | Good fit & continue with other domains |
| Unknown density 11 | I7M688 | 705-796 | I7M688 | 1148-1328 | 15 | 1 | Good fit & continue with other domains |
| Unknown density 12 | I7M688 | 1421-1548 | I7M688 | 1421-1548 | 1 | 1 | Good fit & continue with other domains |

